# Altered primary somatosensory neuron development in a *Pten* heterozygous model for autism spectrum disorder

**DOI:** 10.1101/2023.08.04.552039

**Authors:** Alejandra Fernandez, Nick Sarn, Charis Eng, Kevin M. Wright

## Abstract

Autism spectrum disorder (ASD) is a complex neurodevelopmental disorder characterized by deficits in social interactions, repetitive behaviors, and hyper- or hyposensitivity to sensory stimuli. The mechanisms underlying the emergence of sensory features in ASD are not fully understood, but recent studies in rodent models highlight that these may result from differences in primary sensory neurons themselves. We examined sensory behaviors in a *Pten* haploinsufficient mouse model (*Pten^Het^*) for syndromic ASD and identified elevated responses to mechanical stimuli and a higher threshold to thermal responses. Transcriptomic and *in vivo* anatomical analysis identified alterations in subtype-specific markers of primary somatosensory neurons in *Pten^Het^* dorsal root ganglia (DRG). These defects emerge early during DRG development and involve dysregulation of multiple signaling pathways downstream of *Pten*. Finally, we show that mice harboring an ASD-associated mutation (*Pten^Y69H^*) also show altered expression of somatosensory neuron subtype-specific markers. Together, these results show that precise levels of *Pten* are required for proper somatosensory development and provide insight into the molecular and cellular basis of sensory abnormalities in a model for syndromic ASD.

## Introduction

Altered sensory responsiveness is a defining feature of autism spectrum disorder (ASD) and has been recapitulated in animal models of ASD [1–3]. While much of the research has focused on sensory processing regions of the brain, recent advances show that altered function of primary sensory neurons is frequently seen in models for ASD. Multiple groups have reported alterations in retinal [4], auditory [5], and somatosensory circuits [1–3, 6, 7].

Despite the progress made on characterizing sensory features in animal models of ASD, how these defects emerge during development remains understudied. Somatosensory information is conveyed from the periphery to the CNS by ∼15 molecularly distinct primary neuron subtypes in the dorsal root ganglia (DRG). During embryonic development, nascent DRG neurons delaminate from the neural crest, coalesce into ganglia, and innervate target tissues throughout the body. Neurotrophin signaling through Trk receptors activates multiple signaling pathways that control the survival, differentiation, and maturation of DRG neurons. One of these critical pathways is the PI3K/Akt/mTOR pathway that regulates the survival and differentiation of DRG neurons [8–12]. Phosphatase and tensin homolog (PTEN), is the primary negative regulator of this pathway [13–19]. Heterozygous mutations in *PTEN* have been identified in ∼2% of all ASD cases and represent up to 20% of individuals with ASD and macrocephaly [20]. Sensory phenotypes are a common feature among PTEN-associated ASD [21].

Based on its association with ASD and its critical role in regulating an essential pathway in DRG neuron development, we examined the effects of heterozygous deletion of *Pten* (*Pten^Het^*) on primary somatosensory neurons. We find that loss of a single copy of *Pten* results in alterations in multiple somatosensory behaviors. Bulk RNAseq analysis of adult DRGs from *Pten^Het^* mice indicated shifts in the proportion of neuronal subtypes, which was confirmed by analyzing DRG population markers *in vivo*. We find that these defects manifest at the earliest stages of DRG development and are accompanied by overactivation of both mTOR and GSK3-β signaling cascades downstream of PTEN. In addition, mice harboring a heterozygous ASD-associated mutation that shows predominant nuclear localization (*Pten^Y68H/+^*) phenocopy defects in DRG marker expression [22–24], highlighting the clinical relevance of our model. Taken together, these results show the contribution of PTEN signaling to molecular identity in DRGs, providing insight of the consequences of PTEN mutations and their contribution to sensory features seen in individuals with *PTEN*-related ASD.

## Results

### Alterations in *Pten* leads to somatosensory defects

Germline heterozygous mutations in *PTEN* are associated with ASD and a variety of behavioral phenotypes, including repetitive behaviors, anxiety, social deficits, and sensory features [25–27]. Moreover, a recent study reported alterations in itch and thermal processing following homozygous deletion of *Pten* from adult DRG neurons in mice [28]. However, a close examination of the somatosensory phenotypes caused by germline heterozygous mutations in *Pten* mice remains to be determined. We therefore used a battery of sensory assays broadly focusing on responses to mechanosensation, nociception, and proprioception.

We first assessed mechanosensation by application of an oil droplet on the back of the neck of mice to cause hair deflection. This induces a “wet dog shake (WDS)” response mediated by C-type low threshold mechanoreceptors (C-LTMRs) [29]. In wild type animals (WT), a single oil droplet evoked WDS at a frequency of 11 WDS in a span of 5 minutes, while in *Pten^Het^* mice, the frequency increased by 55% to 16.4 in the same timeframe (**Figure 1A**). We next used an adhesive sticker removal assay, which measures the animal’s ability to localize mechanical stimuli and is thought to engage Aβ- and Aδ-LTMRs [30, 31]. When measuring the latency to remove the sticker, we found that mutant mice displayed a significant delay, with 50% of *Pten^Het^* mice failing to remove the sticker within the 15-minute timeframe. In contrast, only one WT mouse (0.06%) failed to remove the adhesive within the same timeframe (**Figure 1B**). Together, these results demonstrate that *Pten^Het^* mice display alterations in responses to mechanical stimulus in hairy skin.

**Figure 1:**
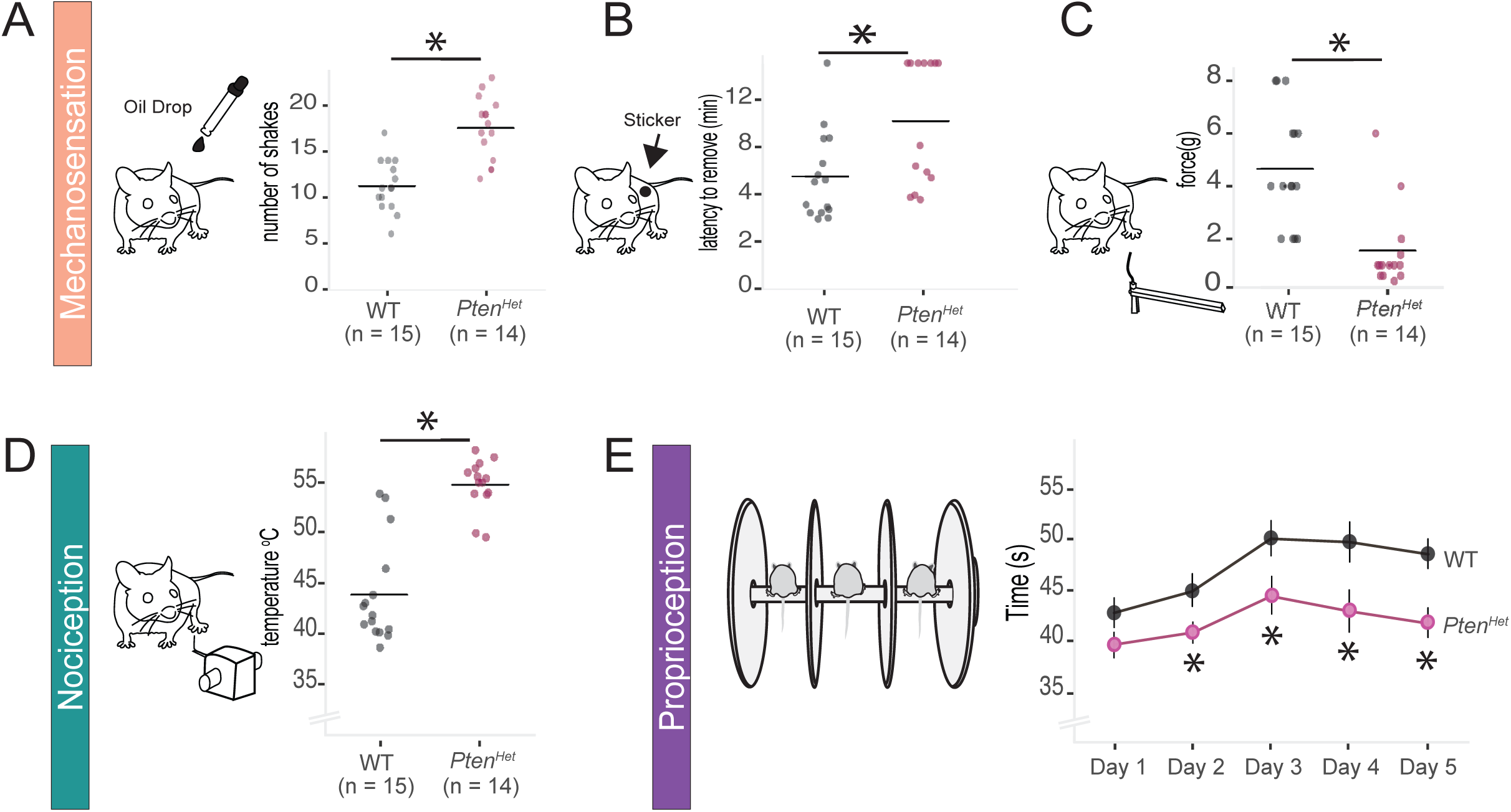
Behavioral abnormalities in different sensory modalities in *Pten^Het^* mutants. **A)** Schematic of oil droplet experiment, in which an oil drop is applied to the back of the neck of a mouse to elicit mechanosensory responses from C-LTMRs (left). Quantification of “wet dog shakes” (WDS) in *Pten* heterozygous mutants and control littermates in a 5-minute interval (right). The average number of responses in mutants is significantly increased by 55% (11.4 ±0.97 WDS in WT vs. 16.4 ±1.28 WDS in *Pten^Het^*, n= 15 WT/ 14 *Pten^Het^*, two-sided unpaired t test p< 0.02). **B)** Illustration of adhesive removal experiment (left) and quantification (right) showing an average increase in latency to remove the adhesive in *Pten^Het^* mutants (5.73 ±0.93 min in WT vs. 10.26 ±1.34 min in *Pten^Het^*, n= 15 WT/ 14 *Pten^Het^*, p< 0.01). 1/15 WT and 7/14 *Pten^Het^* mice failed to remove the adhesive within the 15-minute testing period. **C)** Schematic of Von Frey filament experiment to measure mechanosensory responses to skin indentation in hind paw (left). Heterozygous mutants show lower response threshold to skin indentation (right). Average force needed to elicit a response in mutants was decreased by over 75% in *Pten^Het^* mice (4.6 ±0.57 g in WT vs. 1.54 ±0.18 g in *Pten^Het^*, n= 15 WT/ 14 *Pten^Het^*, p< 0.002). **D)** Schematic of thermal probe test (left) used to detect response to noxious heat (left). *Pten^Het^* mice show increased threshold to thermal stimuli in hind paw when compared to controls (right, 43.82 ±1.30 °C in WT vs. 54.70 ±0.66 °C in *Pten^Het^*, n= 15 WT/ 14 mutants, p< 0.001). **E)** Illustration of accelerating rotarod experiment, where sensory motor control is measured across a span of 5 days (left). Quantification (right) shows mean latency to fall per genotype. Both mutants and controls improve in latency to fall over the first 3 consecutive days of training before stabilizing at a specific level (48 s in WT vs 38 s in *Pten^Het^,* n= 15 WT/ 14 *Pten^Het^*). On days 2-5 there is a significant decrease in the latency to fall in *Pten^Het^* mutants when compared to WT littermates ( day 1: 25.6 ±2.6 s in WT vs 19.3 ±2.2 s in *Pten^Het^*, p = 0.08; day 2: 29.9 ±3.01 s in WT vs 21.9 ± 1.7 s in *Pten^Het^*, p < 0.05; day 3: 40.1 ±3.3 s in WT vs 28.9 ±3.5 s in *Pten^Het^*, p =< 0.05; day 4: 39.5 ±3.5 s in WT vs 25.9 ±4.0 in *Pten^Het^*, p <0.02; day 5: 37.0 ±2.7 s in WT vs 23.5 ±2.7 s in *Pten^Het^*, p<0.002; *Pten^Het^*).

We then assessed mechanical sensitivity in glabrous skin by measuring the threshold to paw withdrawal in response to calibrated Von Frey fibers applied to the plantar surface of the hind paw, which is mediated by Aβ-SA LTMRs [31–33]. We found the threshold of withdrawal responses to be significantly decreased in *Pten^Het^* mice, with filaments ranging between 0.4 to 1.4g eliciting a response, while responses in WT mice were elicited by filaments ranging from 2 to 8g. On average, *Pten^Het^* mice thresholds were reduced by 67% (**Figure 1C**), suggesting mechanosensation hypersensitivity in hind paws.

We then assessed responses to noxious heat stimuli using a thermal probe applied to the plantar surface of the hind paw that heats at a rate of 2.5°C/sec. [33, 34]. WT mice consistently withdrew their paws at temperatures below 50°C. In contrast, *Pten^Het^* mutants exhibited a higher threshold close to 55°C before paw withdrawal (**Figure 1D**). This suggests a higher tolerance to noxious heat in *Pten^Het^* mice.

Finally, we use the accelerating rotarod task to examine proprioception in WT and *Pten^Het^* mice using the accelerating rotarod task. This assay assesses gross motor coordination as well as motor skill learning [35]. We found a significant effect of genotype on rotarod performance on trials conducted on days two through five, with *Pten^Het^* mice showing a reduced latency to fall compared to WT littermates (**Figure 1E**). Taken together, these results provide evidence of the complex alterations in sensory behaviors in mice with a germline deletion of a single copy of *Pten*, as they exhibit hypersensitivity to some stimuli (oil droplet, Von Frey fiber), reduced sensitivity to other stimuli (adhesive sticker, temperature), and impaired sensory-motor behaviors.

### Transcriptional profiles of DRGs are altered in *Pten^Het^* mice

A recent study showed that homozygous deletion of *Pten* from adult DRGs results in altered itch behavior due to the upregulation of multiple itch-related genes [28]. To more comprehensively understand how heterozygous deletion of *Pten* affects gene expression in DRGs, we performed bulk RNA-sequencing (RNA-seq) of whole DRGs isolated from adult *Pten^Het^* and WT littermate control mice (**Figure 2A**). We identified 285 differentially expressed genes (DEG) in *Pten^Het^* mutant mice relative to WT, with 41 genes upregulated and 244 genes downregulated (**Figure 2B and Table S1**). Overall gene expression changes were relatively small, with most of the DEG either up- or downregulated by <1.5 Log2 fold change (Log2FC) (**Figure 2C**). The widespread yet relatively small change in gene expression is consistent with studies that have examined the effect of single copy deletion or mutation of *Pten* in other neuronal populations [24, 36]. Gene Ontology (GO) analysis of DEGs identified significant changes to gene networks involved in metabolism, RNA processing, signal transduction, and the ubiquitination pathway (**Figure 2D**). Surprisingly, *Pten* was slightly upregulated in mutant DRG samples, potentially due to epigenetic factors or altered distribution of *Pten* expression throughout DRG subtypes [37].

**Figure 2:**
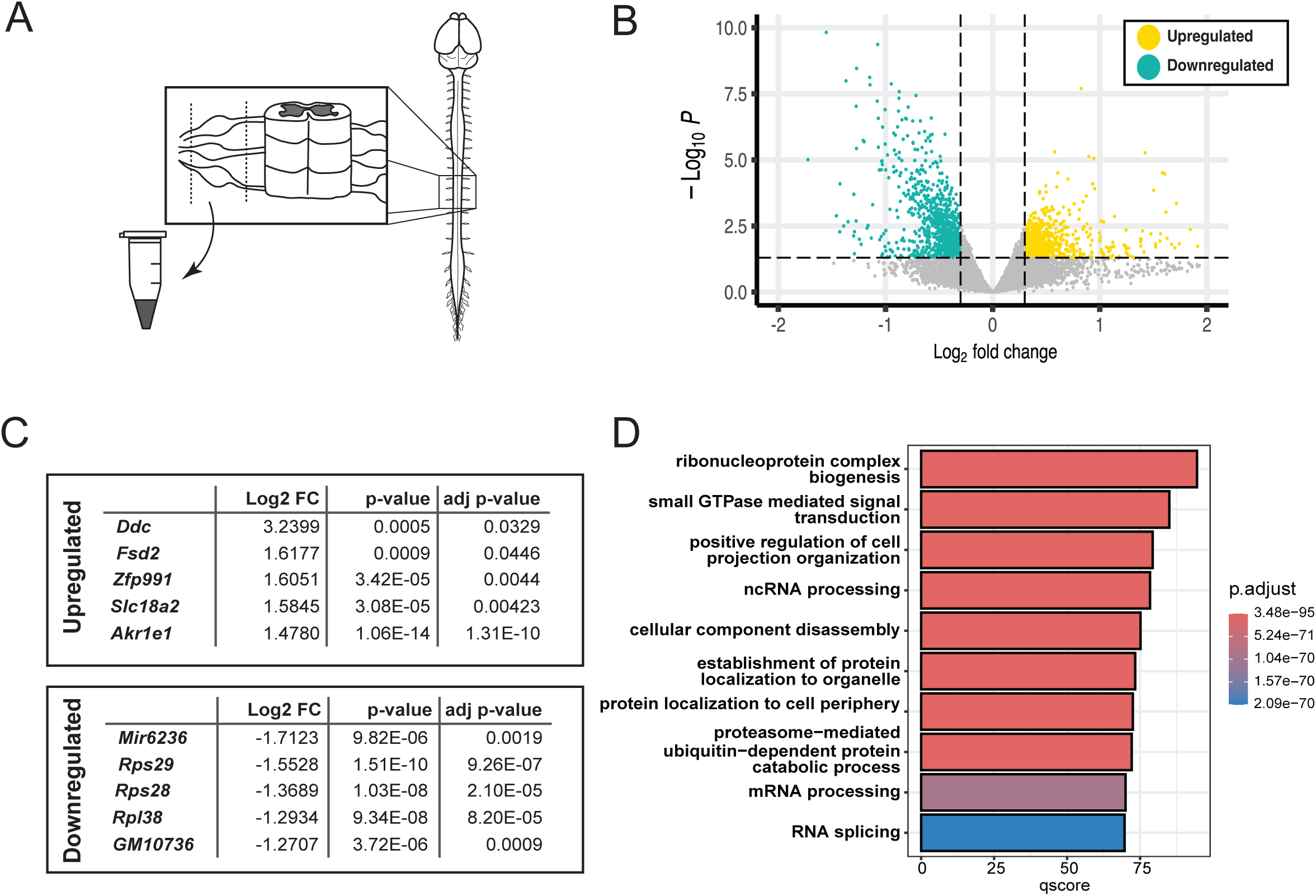
Altered transcriptome in DRGs from *Pten^Het^* mutants. **A)** Schematic of experimental design. Whole DRGs along the entire rostro-causal axis were dissected from *Pten^Het^* mutants and WT littermates and processed for bulk RNA sequencing (RNAseq). **B)** Volcano plot highlighting genes that are significantly upregulated (41 genes, LFC > 0.3, yellow, right) and downregulated (244 genes, LFC < -0.3, turquoise, left) in *Pten^Het^* mutants (adjusted p value < 0.05). **C)** List of top differentially expressed genes (top 5 upregulated genes on top, top 5 downregulated genes bottom). **D)** Barplot showing Gene Ontology analysis for differentially expressed genes. Bars are color-coded from blue to red based on the adjusted p-value.

We then examined whether loss of a single copy of *Pten* affected transcriptional control of specific DRG subtypes by focusing on genes enriched in each somatosensory neuron subtype as defined by an available single-cell RNA seq dataset [38]. We examined the top 50 enriched genes within each subtype and identified those that were significantly altered in *Pten^Het^* mutants (**Figure 3A**). We found that about ∼10% of the subtype-specific markers for each neuronal subtype were differentially expressed in *Pten^Het^* DRGs (**Figure 3B**). We used DeconV, a deconvolution algorithm that leverages gene expression profiles from a reference sample along with probability distributions, to infer the proportional abundance of distinct cell types within a target bulk RNA-seq sample [39] (**Figure 3C**). This approach detected a potentially altered distribution of DRG subtype transcriptomes in *Pten^Het^* mutants when compared to controls, particularly in populations of C-LTMRs, peptidergic nociceptors, and proprioceptors (**Figure 3D**).

**Figure 3:**
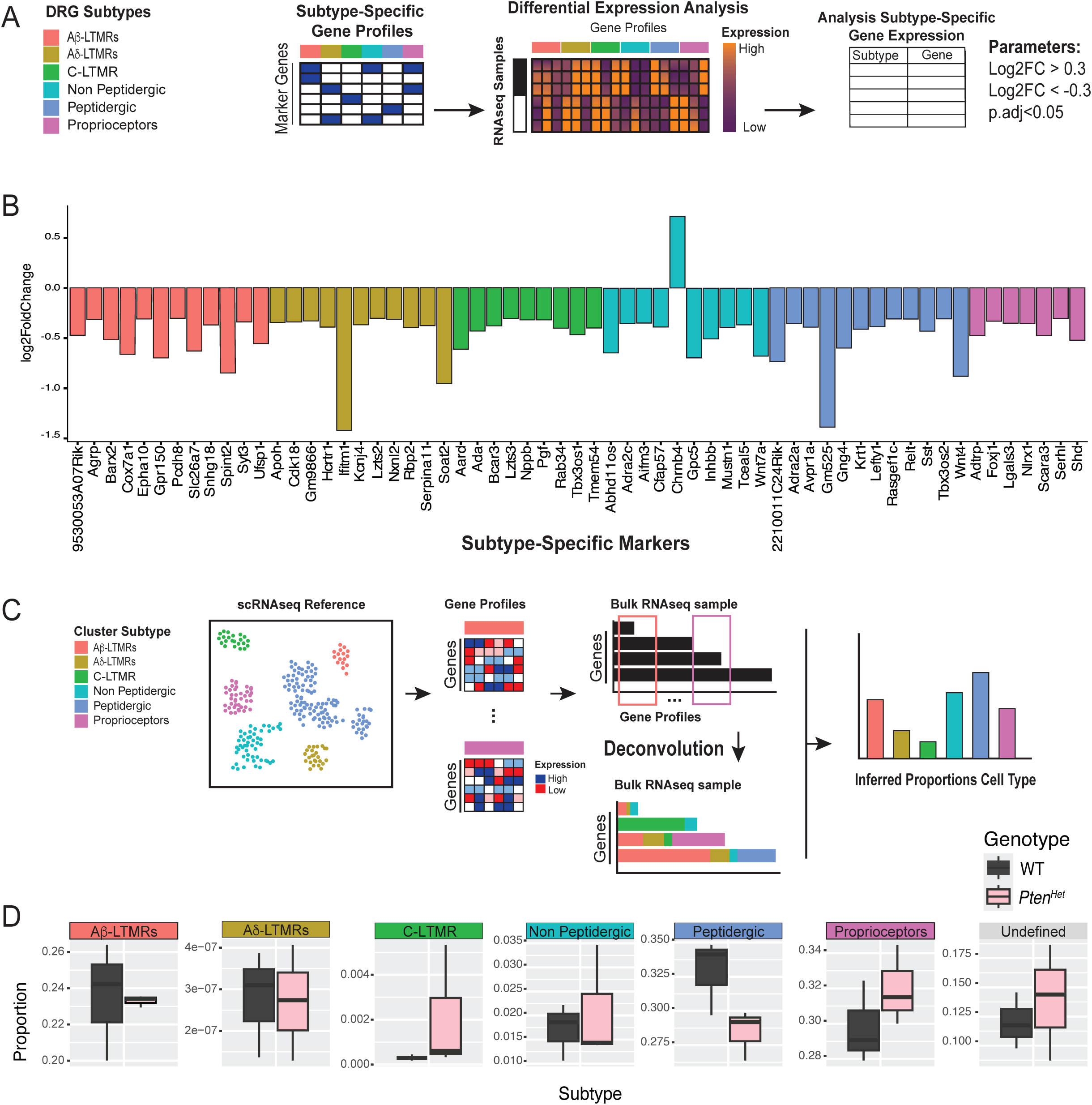
Alterations in DRG neuron subtype-specific genes in *Pten^Het^* mutants. **A)** Schematic experimental workflow to examine differential expression of DRG subtype-specific genes in RNA-seq data from *Pten^Het^* and WT littermate mice. Analysis is focused on the major neuronal subpopulations in DRG (left), with multiple subtypes of peptidergic and non-peptidergic nociceptors combined into two clusters. Subtype-specific gene profiles were determined based on DRG subtype-specific markers from a scRNA-seq dataset [38]. The top 50 differentially expressed genes within each of the subtype-specific profiles were analyzed. **B).** Bar plot showing subtype-specific genes with differential expression between *Pten^Het^* and WT littermate mice. Bar colors correspond to each DRG population subtype (adjusted p< 0.05). **C)** Computational framework to infer proportions of DRG subtypes from RNA-seq data using DeconV [39]. scRNAseq data containing expression profiles from each DRG population subtype is used as a reference dataset (left). Gene profiles for each population are extracted and the probabilistic framework of DeconV is used to estimate proportions of each subtype, based on an assumed linear-sum-property between single-cell and bulk gene expression (middle). DeconV then provides the proportions of each subtype as an output (right). **D)** Inferred quantification of DRG population subtypes in WT and *Pten^Het^* by probabilistic cell type deconvolution. Quantification suggests alterations to distribution of C-LTMRs, peptidergic nociceptors, and proprioceptor populations in *Pten^Het^* mutants.

### DRG subtype-specific marker expression is altered in *Pten^Het^* mice

Based on the DeconV prediction of altered DRG subtype transcriptomes in *Pten^Het^* mutants, we evaluated DRG subtype populations *in vivo*. We collected hindlimb-innervating DRGs (lumbar level 3-4; L3-L4) from adult *Pten^Het^* mutants and WT control littermates and quantified the various DRG populations using subtype-specific markers (**Figure 4A**). We found no difference in the total number of Isl1/2^+^ DRG neurons between mutants and controls (**Figure 4B**). In contrast, we identified alterations in the proportion of DRG neuron subtype-specific markers in *Pten^Het^* mutants, with some populations affected while others were spared. There was a 156% increase in Tyrosine Hydroxylase positive (TH^+^) C-LTMRs and a 56% increase in TrkC^+^ proprioceptors in *Pten^Het^* mutants. TrkA^+^ peptidergic nociceptors were reduced by 27% in *Pten^Het^* mutants relative to WT littermates, whereas IB4^+^ non-peptidergic nociceptors were unaffected.

**Figure 4:**
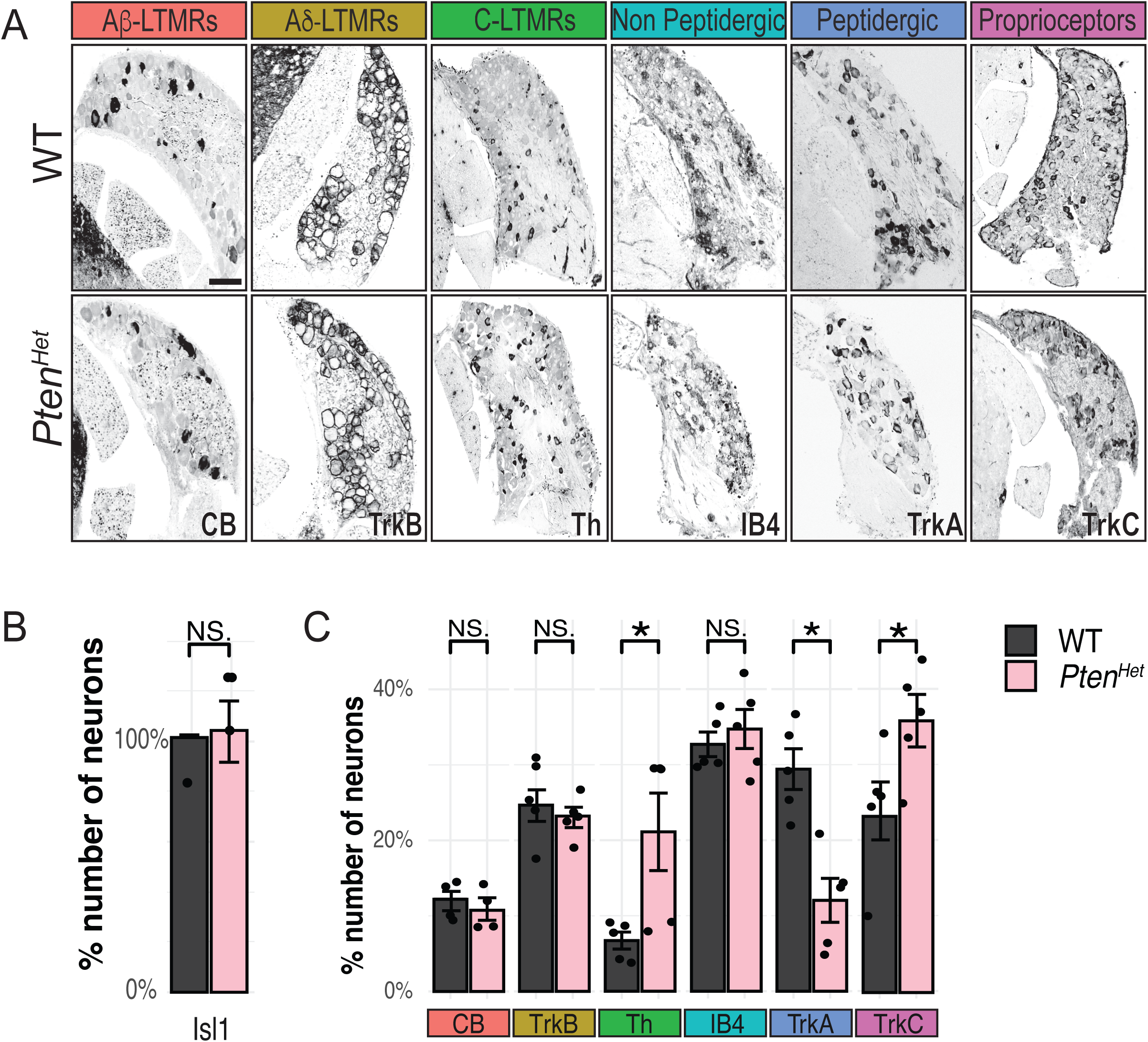
Expression of subtype-specific markers is altered in *Pten^Het^* mutants. **A)** Representative images of cryosections of WT and *Pten^Het^* adult DRGs with specific markers (Scale 100 µm). **B)** Quantification of the proportion of Isl1/2^+^ neurons shows no significant difference in the total population of neurons in *Pten^Het^* and WT littermate DRGs (n=3, p=0.57, Student’s t-test). **C)** Quantification of each major DRG neuronal subtype in WT and *Pten^Het^* mutants. There were no significant changes in the number of Calbindin positive (CB^+^) Aβ-LTMRs or TrkB^+^ Aδ-LTMRs (CB^+^ Aβ-LTMR: 11.5 ±1.1% in WT vs. 10.4 ±1.3% in *Pten^Het^*, n=5, p=0.23 and TrkB^+^ 25.5±2.0% in WT vs. 23.2 ± 1.4% in *Pten^Het^*, n=5, p=0.21, Student’s t-test). Tyrosine hydroxylase positive (TH^+^) C-LTMRs are increased in *Pten^Het^* mutants compared to WT littermates (6.7 ±1.1% in WT vs. 21.1 ±5.1% in *Pten^Het^*, n=5, p<0.01, Student’s t-test). Quantification of IB4-positive non-peptidergic nociceptors shows no change in *Pten^Het^* mutants compared to WT controls (32.6 ± 1.6% in WT vs. 34.7±2.5% in *Pten^Het^*, n=5, p=0.26, Student’s t-test). TrkA positive peptidergic nociceptors are decreased in *Pten^Het^* mutants (29.4 ±2.6% in WT vs. 12.0 ±2.9% in *Pten^Het^*, n=5, p<0.002, Student’s t-test). TrkC positive (TrkC^+^) proprioceptors were increased in *Pten^Het^* mutants compared to WT littermates (24.4± 3.9% in WT vs. 37.0 ±3.5% in *Pten^Het^*, n=5 p<0.02, Student’s t-test).

On the other hand, Calbindin positive (CB^+^) Aβ-LTMRs and TrkB^+^ Aδ-LTMRs were unaffected in *Pten^Het^* mutants (**Figure 4A, C**). These results were consistent with the predictions from the transcriptomic data, and results suggest that loss of a single copy of *Pten* causes alterations in DRG subtype-specific marker expression and potential defects in subtype-specific diversification.

### Early DRG development is abnormal in *Pten^Het^* mutants

DRG subtype diversification begins embryonically and ultimately generates the full complement of genetically and physiologically distinct subtypes of somatosensory neurons (**Figure 5A**). Large/medium diameter neurons expressing TrkC and/or TrkB are born first and give rise to most myelinated neurons including proprioceptors and mechanoreceptors. TrkA expressing small diameter neurons are born shortly after the TrkC/TrkB populations and differentiate into unmyelinated nociceptors and C-fiber LTMRs [19]. We examined proportions of TrkA, TrkB, and TrkC positive neurons between E10.5 and E14.5 to determine how early differentiation was affected in *Pten^Het^* mutants. We found a significant decrease in the number of TrkA^+^ neurons and a significant increase in the number of TrkC^+^ neurons in *Pten^Het^* mice compared to WT littermates at all ages examined (**Figure 5B, D**). In contrast, the number of TrkB^+^ neurons between E10.5-E14.5 was unaffected, in line with our results in adult mice (**Figure 5C**). We also examined whether changes in cell proliferation or developmental apoptosis contributed to the alterations in DRG subtype proportions using EdU birth dating and activated Caspase-3 staining, respectively, and we observed no significant differences in neuronal progenitor proliferation or developmental apoptosis between *Pten^Het^* mutants and WT littermates in developing DRGs (**Supplemental Figure 1**).

**Figure 5:**
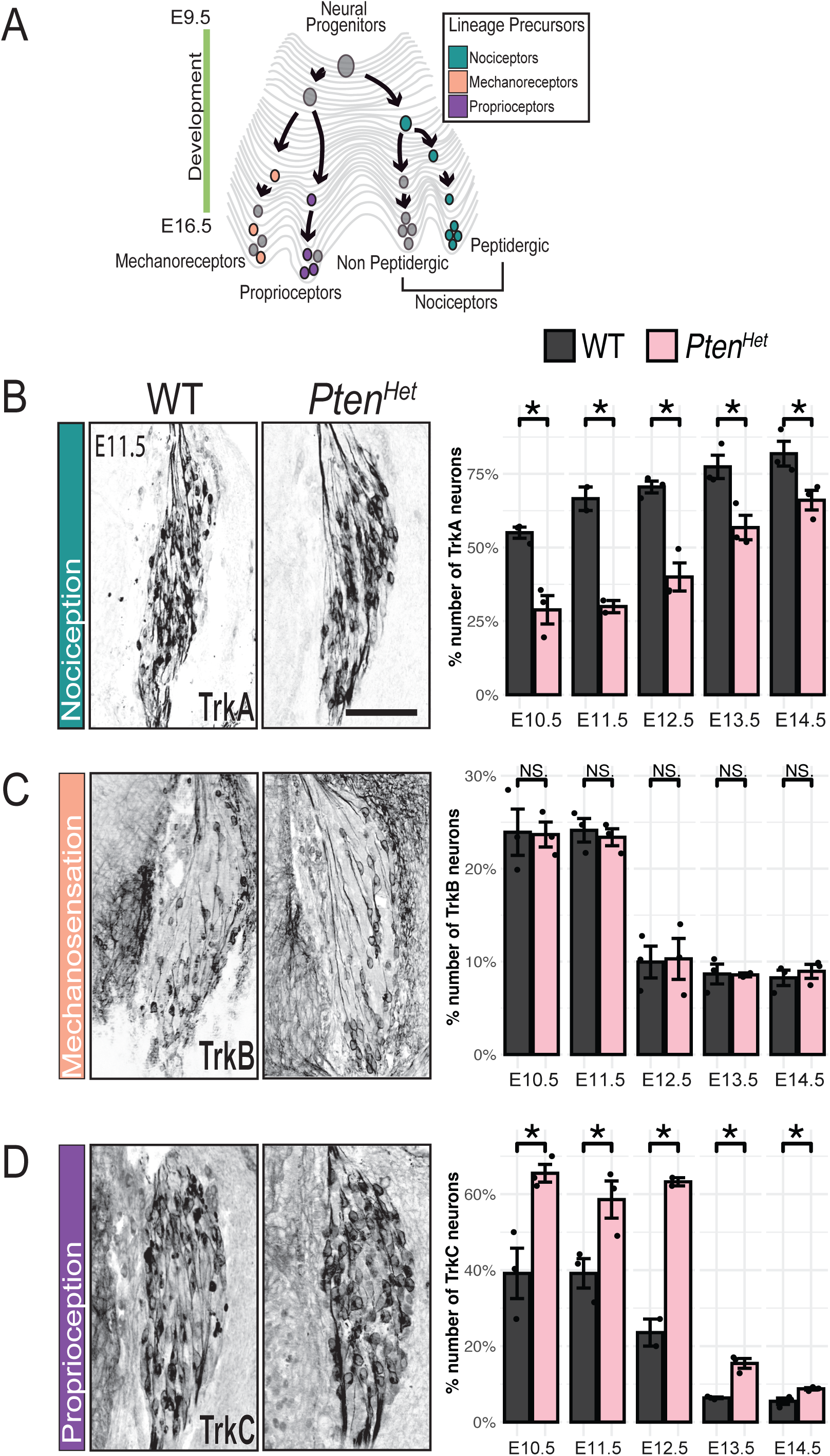
Neuronal differentiation is altered in *Pten^Het^* mutants early in development. **A)** Illustration of developmental landscape showing progression of DRG neuron diversification. **B-D)** Representative images of cryosections of WT and *Pten^Het^* E11.5 DRGs with markers of each subtype (left panels) and quantification (right panels) (Scale 100 µm). **B)** Representative images of DRGs labeled with TrkA (left), quantification (right) shows a decrease in the number of TrkA positive neurons in *Pten^Het^* mutants compared to WT littermates throughout development (n=3, E10.5: p<0.004; E11.5: p<0.03; E12.5: p<0.02, E13.5: p<0.045, E14.5: p<0.03, Student’s t-test). **C)** Images showing DRGs labeled with TrkB (left), quantification (right) shows no changes the number of TrkB^+^ neurons in *Pten^Het^* mutants (n=3, E10.5: p=0.46; E11.5: p=0.32; E12.5: p=0.45; E13.5: p=0.47; E14.5: p=0.28, Student’s t-test). **D)** Images of E11.5 DRGs labeled with TrkC (left), quantification (right) shows an increase in the proportion of TrkC+ neurons in *Pten^Het^* mutants compared to WT littermates after E10.5 (E10.5: p<0.05; E11.5 p<0.05; E12.5: p<0.02; E13.5: p<0.05; E14.5: p<0.01, Student’s t-test).

These results suggest that appropriate levels of Pten are critical for the proper differentiation of DRG subtypes.

### Downstream signaling pathways are altered in *Pten^Het^* mutants

Given that *Pten* haploinsufficiency has distinct effects on developing DRG subtypes, we reasoned that signaling cascades downstream of PTEN could potentially be differentially impacted in population subtypes. PTEN is the primary negative regulator of PI3K/AKT signaling and modulates activity of both the TSC/mTOR/S6 pathway and the GSK-3β/β-catenin pathway [40–42] (**Figure 6A**). We examined the proportions of DRG neurons engaged in each signaling cascade at E11.5, using phosphoS6 (pS6) immunostaining as a readout of TSC/mTOR pathway activity. We used a *TCF/Lef:H2B-GFP* reporter mouse that provides a readout of WNT/β-catenin signaling activity to examine GSK-3β/β-catenin activation [43]. We found an increased proportion of Isl1^+^ neurons that were positive for both pS6 and GFP in *Pten^Het^* mutants when compared to WT (**Figure 6B, C**). Closer examination of each DRG subtype revealed population-specific activation of PTEN-dependent downstream signaling. Both TrkA^+^ and TrkC^+^ neuronal subtypes showed a significant increase in the proportion of neurons positive for pS6, while there was no change in the proportion of pS6^+^ TrkB^+^ neurons (**Figure 6D-G**). In contrast, only the TrkC^+^ population had a significantly increased proportion of GFP^+^ neurons indicative of Gsk-3β/β-catenin activation. These results provide evidence for subtype-specific alterations of signaling cascades due to *Pten* haploinsufficiency during DRG development.

**Figure 6:**
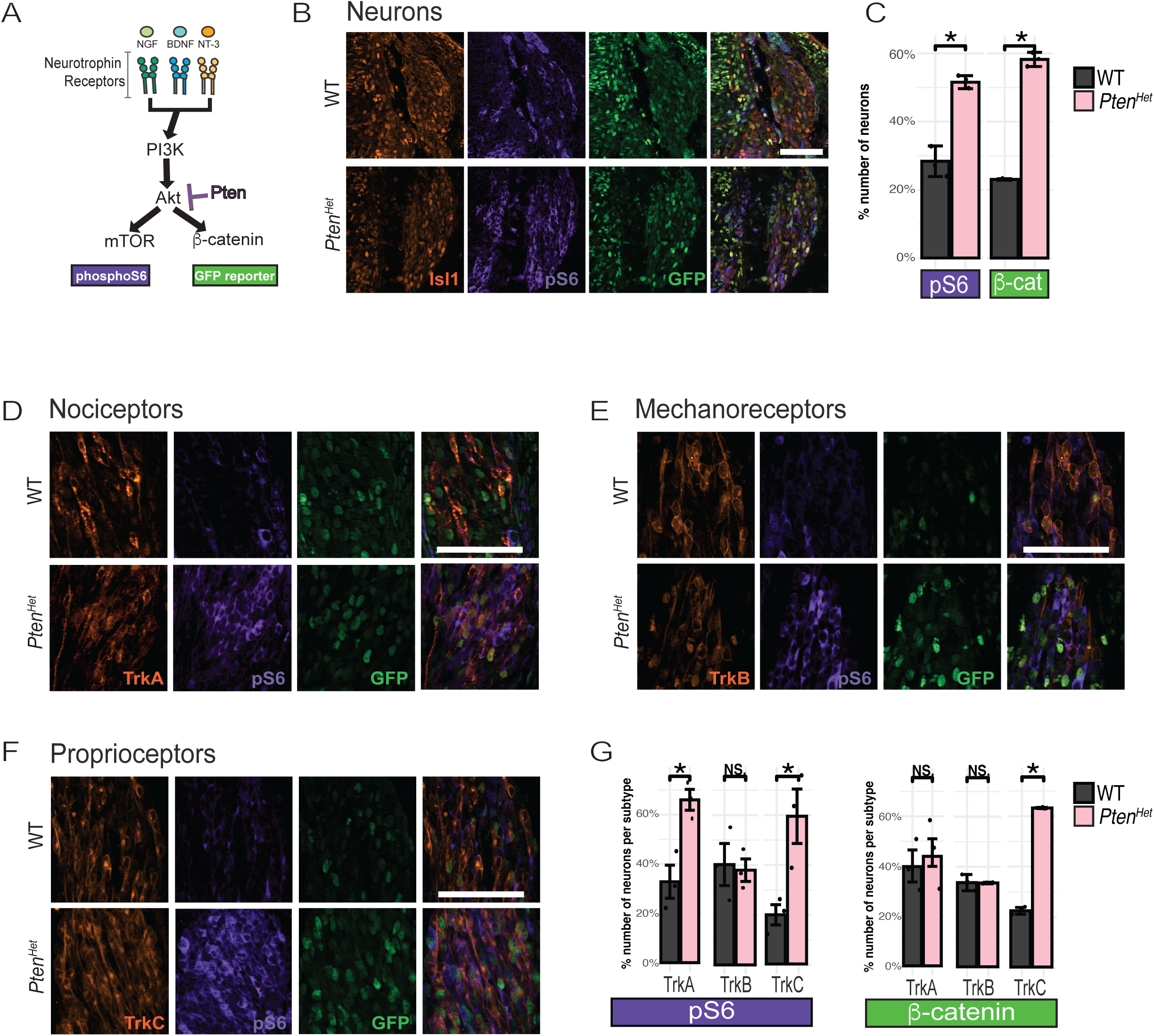
P*t*en haploinsufficiency alters signaling downstream of PI3K/AKT in TrkA and TrkC lineages. **A)** Schematic representation of mTOR and GSK-3β signaling pathways downstream of Trk receptor activation of PI3K/AKT. **B, D-F)** Representative images of E11.5 DRGs of WT and *Pten^Het^* littermates immunolabeled with Isl1 (B), TrkA (D), TrkB (E), TrkC (F) (orange), phospho-S6 (pS6) (purple) and GFP (green) to measure activity of the mTOR and GSK-3β/β-catenin pathways, respectively (Scale 100 µm). **C)** Quantification of Isl1^+^ neurons positive for activation of pS6 (left) and β-catenin (β-cat, right) in E11.5 DRGs from WT and *Pten^Het^* mice. There was an increase in the proportion of pS6^+^/Isl1^+^ neurons in *Pten^Het^* compared to WT (28.3±4.4% in WT vs. 51.5 ±1.9 in *Pten^Het^*, n=3, p<0.041, Student’s t-test). The proportion of GFP^+^/Isl1^+^ neurons was also increased in mutants compared to controls (23.0±0.2% in WT vs. 58.2 ±2.0 in *Pten^Het^*, <u>n</u>=3, p<0.005, Student’s t-test). **G)** Quantification of pS6 (left) and GFP (right) positive neurons within each DRG neuron subtype. Within the TrkA population, the pS6^+^/TrkA^+^ neurons were increased in *Pten^Het^* compared to WT littermates (left, 32.8±8.2% in WT vs. 66.0 ±5.1 in *Pten^Het^*, n=3, p<0.02, Student’s t-test), while there was no significant change in the number of GFP^+^/TrkA^+^ neurons (39.7±6.9% in WT vs. 45.2 ±5.5 in *Pten^Het^*, n=3, p=0.30, Student’s t-test). Quantification in TrkB^+^ neurons shows no significant change in the proportions of pS6^+^/TrkB^+^ neurons between *Pten^Het^* and WT controls (39.2±14.2% in WT vs. 37.0 ±2.8 in *Pten^Het^*, n=3, p=0.41, Student’s t-test, left) or in the proportion of GFP^+^/TrkB^+^ neurons (38.4±3.7% in WT vs. 38.2 ±0.3 in *Pten^Het^*, p=0.96, Student’s t-test). There was an increased number of pS6^+^/TrkC^+^ neurons in *Pten^Het^* compared to WT littermates (19.3±6.8% in WT vs. 58.2 ±13.1 in *Pten^Het^*, n=3, p<0.03, Student’s t-test, left), and GFP^+^/TrkC^+^ neurons in *Pten^Het^* compared to controls (25.2±1.5% in WT vs. 71.0 ±0.2 in *Pten^Het^*, n=3, p<0.002, Student’s t-test).

### DRG neuron diversification is abnormal in mice with an ASD-associated mutation in *Pten*

Given the prevalence of *PTEN* mutations within the ASD population and the high incidence of sensory phenotypes in ASD [44, 45], we examined DRG subtype markers in a mouse model carrying a patient-specific *PTEN* mutation (*Pten^Y68H/+^*). This mutation affects the phosphatase domain and results in reduced protein stability, phosphatase activity, and predominantly nuclear localization in CNS neurons [24, 46, 47]. We found that *Pten^Y68H/+^* mice have alterations in DRG subtype proportions that mirror our results in *Pten^Het^* mutants. There was a 79% increase in TH^+^ C-LTMRs and 96% increase in TrkC^+^ neurons in *Pten^Y68H/+^* mice when compared to WT littermates. The proportion of TrkA^+^ peptidergic nociceptors was reduced by 50%, whereas the population of IB4^+^ non-peptidergic nociceptors was unaffected. In contrast, there was no change in the proportion of CB^+^ Aβ LTMRs or TrkB^+^ Aδ-LTMRs (**Figure 7A, B**). The nearly identical phenotypes of *Pten^Y68H/+^* mice and *Pten^Het^* mice highlights the clinical relevance of our findings that identify a critical role for precise Pten levels in regulating DRG development and the expression of subtype-specific markers.

**Figure 7:**
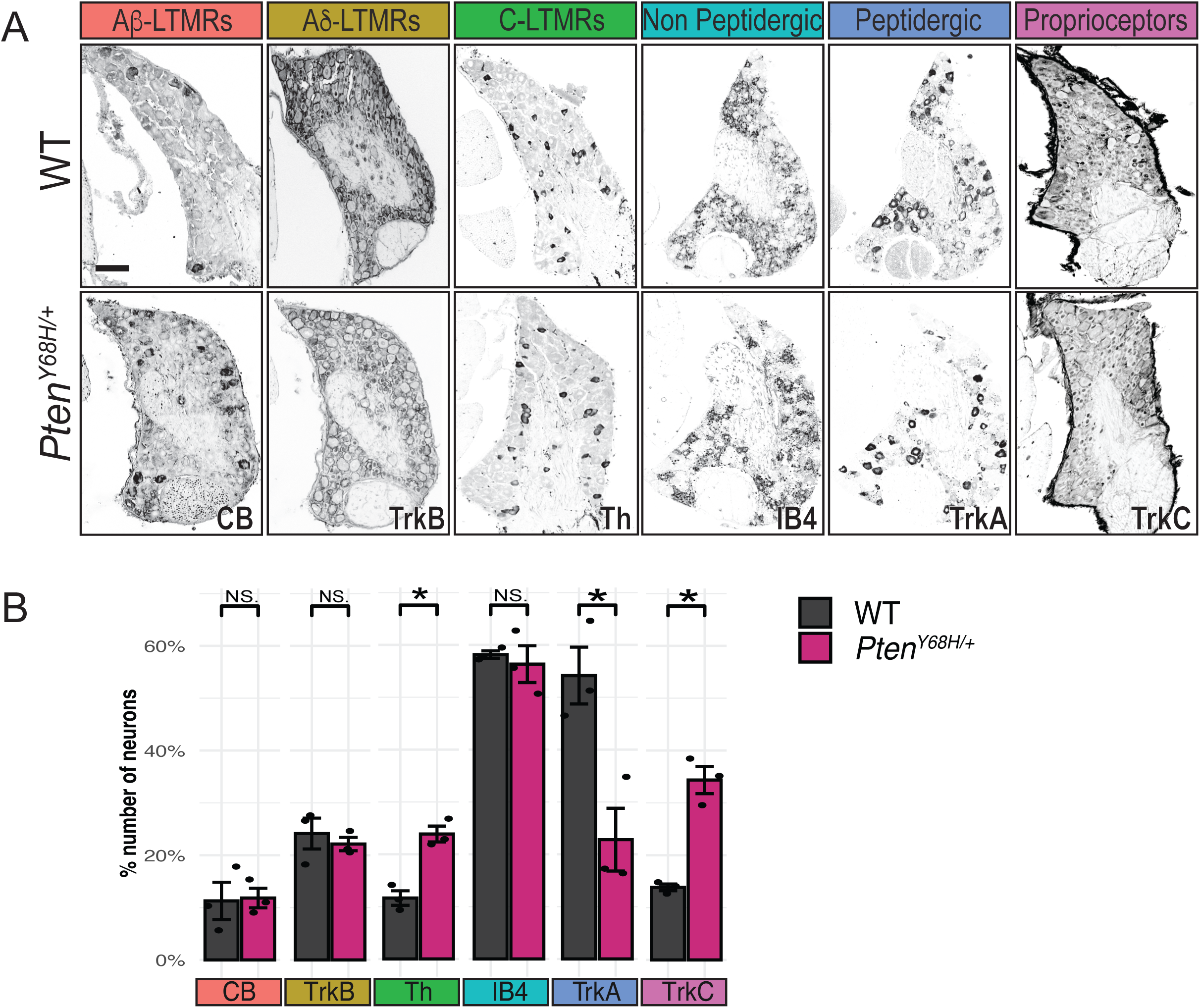
DRG neuron diversification is abnormal in *Pten^Y68H/+^* mice. **A)** Representative images of cryosections from WT (top) and *Pten^Y68H/+^* (bottom) P21 DRGs labeled with specific markers for select DRG subtypes (Scale 100 µm). **B)** Quantification shows subtype-specific alterations in DRG neurons in *Pten^Y68H/+^* compared to WT littermates. The proportions of CB^+^ Aβ-LTMRs and TrkB^+^ Aδ-LTMR were unchanged in *Pten^Y68H/+^* mutants, while there was a significant increase in the proportion of TH^+^ c-LTMRs (CB1: 11.2 ±3.5% in WT vs. 11.8± 1.8% in *Pten^Y68H/+^*, n=3 p=0.45; TrkB: 24.1±2.9% in WT vs. 22.1±1.2% in *Pten^Y68H/+^*, n=3 p=0.28; TH: 11.6±1.3% in WT vs. 23.8±1.5% in *Pten^Y68H/+^*, n=3 p<0.002). There were no changes in the proportion of IB4^+^ non-peptidergic nociceptors and a significant decrease in the proportion of TrkA^+^ peptidergic nociceptors in *Pten^Y68H/+^* when compared to littermate controls (IB4: 58.1±6.9% in WT vs. 56.3±3.5% in *Pten^Y68H/+^*, n=3 p=0.31; TrkA: 54.1±5.4% in WT vs. 22.7±6.0 in *Pten^Y68H/+^*, n=3 p<0.01, Student’s t-test). There was a significant increase in the proportion of TrkC^+^ neurons in *Pten^Y68H/+^* mutants compared to WT littermate controls (13.8±0.6% in WT vs. 34.3±2.6% in *Pten^Y68H/+^*, n=3 p<0.001).

## Discussion

PTEN is the primary negative regulator of PI3K/AKT signaling and has been extensively studied for its role in the development, function, and regeneration of the nervous system. However, little is known about its contribution to the *in vivo* development and function of peripheral somatosensory neurons in the DRG. Using a combination of behavioral analysis, transcriptional profiling, and *in vivo* analysis of DRG development, we have identified a critical role for *Pten* in the control of subtype-specific gene profiles in primary sensory neurons. These results provide a molecular basis for developmental mechanisms underlying the emergence of complex sensory alterations in a mouse model for ASD.

*PTEN* plays a critical role in many neurodevelopmental processes [48]. *Pten* heterozygous mouse mutants recapitulate many of the behavioral phenotypes seen in patients with ASD, including social deficits, repetitive behaviors, and anxiety. At the cellular level, this is accompanied by widespread brain overgrowth, altered scaling of neuronal and glial cell populations, increased axonal branching, hyperconnectivity, and decreased intrinsic excitability due to altered expression of ion channels [42, 49–51].

These results indicate multiple distinct mechanisms underlying the complexity of behavioral phenotypes in *Pten* heterozygous models [42, 51]. A recent transcriptomic analysis revealed that during cortical development, heterozygous deletion of *Pten* affects downstream signaling differently in astrocytes and neurons [36]. Why some cell types are more sensitive to *Pten* gene dosage than others remains a subject of current investigation.

Our transcriptomic data (**Figure 3**) and *in vivo* analysis (**Figure 4, 7**) provide evidence supporting a role for proper PTEN levels in DRG neuron subtype differentiation. Notably, we see the same defects in DRG neuron diversification in both *Pten^Het^* and *Pten^Y68H/+^* mice. We show that these defects arise early in development (**Figure 5**) and appear to engage canonical pathways downstream of PTEN (**Figure 6**). This is supported by the observation that both TSC/mTOR and GSK-3β/β-Catenin signaling pathways are dysregulated in *Pten^Het^* DRGs. Interestingly, regulation of these pathways seems to be specifically regulated in different DRG populations. In *Pten^Het^* mutants, both mTOR and GSK-3β/β-Catenin pathways are hyperactive in TrkC^+^ DRG neurons in *Pten^Het^* mutants, whereas only pS6 was significantly elevated in TrkA^+^ DRG neurons and both pathways appeared normal in TrkB^+^ DRG neurons. This could reflect intrinsic differences in how signaling pathways downstream of specific Trk receptors are regulated, providing insight into why some DRG subtypes are more sensitive to heterozygous deletion of *Pten*. Mutations in both TSC1/2 and β-catenin are associated with ASD, but the interplay between signaling dynamics of these pathways and DRG cell-fate specification remain to be determined [52, 53].

Somatosensory phenotypes in *Pten^Het^* mutants shed light into the consequences of DRG developmental defects due to *Pten* signaling. Interestingly, we found a correlation between behavioral abnormalities and alterations in specific DRG subtype proportions. The increased frequency of wet dog shakes in *Pten^Het^* mice in the oil drop test (**Figure 1A**) may be due to the increased density of TH^+^ C-LTMRs (**Figure 4A, C**). It is possible that an increase in C-LTMR numbers in DRGs may be accompanied by increased receptive field coverage in hairy skin that causes light touch hypersensitivity [54]. On the other hand, deficits in sensorimotor control (**Figure 1E**) correlate to the increase in TrkC^+^ proprioceptors in *Pten^Het^* mutants. Proprioceptive sensory feedback is required for the proper development of motor and sensory neuronal circuits and coordination of locomotor activity [55, 56]. Therefore, it is possible that overrepresentation of proprioceptors alters the precision of sensorimotor control in *Pten^Het^* mutants. Given the complex representation of pain responses across many subtypes of multimodal nociceptors, the relationship between the higher temperature threshold and underrepresentation of TrkA^+^ neurons in *Pten^Het^* mutants is unclear. Importantly, at this point we cannot distinguish whether the changes in adult neuron subtype markers are entirely due to altered populations of DRG subtypes, or due to ectopic expression, misexpression, or reduced labeling from individual markers. However, our embryonic data showing early changes in the populations of TrkA^+^ and TrkC^+^ DRG lineages suggest a phenotype encompassing alterations to the genetic profiles of each subtype of primary sensory neurons.

Our results support recent findings from rodent models showing that mutations in ASD-susceptibility genes affect sensory circuit development and/or function [2, 6, 57–62]. Rats with mutations in *Mecp2* show abnormal mechanical and thermal sensitivity due to abnormal sensory neuron postnatal development and function [7]. Moreover, deletion of *Mecp2* or *Gabrb3* in somatosensory neurons results in tactile deficits, defects in social interactions, and anxiety-like behaviors, all of which are associated with

ASD [2]. Interestingly, the developmental timing of somatosensory dysfunction and emergence of ASD-like behaviors in rodent models is tightly linked [3]. Our results in this study reinforce the developmental origin of sensory defects in models for ASD. Pharmacologically reducing tactile overactivity in *Mecp2* and *Shank3* mutants reduced anxiety-like behaviors and some social deficits [6], suggesting that sensory contributions to ASD behaviors may be amenable to therapeutic intervention. Considering that up to 90 percent of individuals with ASD experience some form of atypical sensory sensitivity [45], additional research is needed to better understand the molecular and cellular basis for these phenotypes.

## Acknowledgements

We would like to thank the members of the Wright laboratory for their assistance and discussion throughout the course of this study. Special thanks to Dr. Matthew Pomaville for his insight and help with experimental troubleshooting. We also thank Dr. Martin Riccomagno, Dr. Brian O’Roak, and Dr. Arpiar Saunders for their helpful suggestions on the manuscript. This work was supported by NIH grant R01NS091027 (K.M.W.), OHSU Fellowship for Diversity in Research (A.F.), Collins Medical Trust (A.F.) and NIH grant K01NS116168 (A.F.). C.E. passed away during the preparation of this manuscript and was the Sondra J and Stephen R Hardis Chair of Cancer Genomic Medicine at the Cleveland Clinic and thanks the Ambrose Monell Foundation.

## Materials and Methods

### Mouse lines

All experiments were performed in accordance with OHSU IACUC Protocol #IS00000539. Animals were housed in standard conditions and cared for by the Department of Comparative Medicine at Oregon Health & Science University. Mice were maintained in a 12-hour light/dark cycle and provided food and water *ad libitum*. For experiments involving embryonic timepoints, noon on the date that a vaginal plug was detected was considered E0.5. For experiments involving constitutive *Pten* heterozygous mice, male *Pten^Het^* breeders (*B6;129P2-Pten^tm1Mak^/Mmjax*, MMRRC stock #42059 [63]) were crossed with CD1 females to generate experimental mice on a mixed genetic background. For examining phenotypes related to the patient-specific Y68H *Pten* mutation, *Pten^Y68H/^*^+^ male breeders [24] were crossed to CD1 females to generate experimental mice. We examined activation of the GSK-3β/β-Catenin pathway, using *Pten^Het^* breeders crossed to a *TCF/Lef:H2B-EGFP* reporter line (*Tg(TCF/Lef1-HIST1H2BB/EGFP)61Hadj/J*, MMRRC stock #013752 [43]). All animals were group housed with littermates and maintained on a mixed genetic background. Mice of both sexes were used for all experiments, and WT littermates were used as controls.

### Behavioral Assays

#### Oil drop test

Test was performed as previously described [29]. In brief, animals were placed in a plexiglass box where they were allowed to acclimate and explore for 15 minutes. After the acclimation period, we delivered a 50µl drop of sunflower oil (stabilized at room temperature) and observed behaviors for 5 minutes.

#### Adhesive removal test

Light touch was assessed as previously described [30]. Mice were placed in a plexiglass container, and after a period of acclamation of at least 5

minutes, a small piece of tape (7mm diameter) was carefully placed onto the back of the coat of an unrestrained mouse. The animal was observed for 15 minutes, and the number of responses to the tape stimulus were recorded.

#### Von Frey assay

We used a Dynamic Plantar Aesthesiometer (UgoBasile), which has calibrated von Frey fibers and a movable force-actuator, to measure mechanical somatosensation. Animals were enclosed but unrestrained in an acrylic chamber placed over a meshed floor during the duration of the experiments. The actuator was placed below the paw of the animal to confer different degrees of force *via* a von Frey filament, starting below threshold of detection and increasing until the paw withdrawal reflex was elicited.

#### Heat nociception

Animals were placed in an acrylic box over a grated platform unrestrained during the duration of the experiment. After a 5-minute acclimation period, a blunt probe with a force of 1g was applied to the plantar surface to the animal’s hindlimb. The metal probe touched the paw, heating up at a rate of 2.5°C/sec to a maximum temperature of 60°C. Animals were monitored and their reaction times and temperature at the time of withdrawal were recorded. Testing was performed 3 times per animal, with a 60 second recovery time between intervals.

#### Tissue preparation and RNA isolation

For transcriptome profiling by RNA sequencing, whole DRGs along the entire rostro-caudal axis from 4 animals of each genotype were dissected, collected in 1.5 mL Eppendorf tubes, pelleted and snap-frozen in dry ice. Frozen tissue was shipped on dry ice to Genewiz (Azenta Life Sciences) for RNA sequencing. RNA sequencing workflow included RNA isolation, PolyA selection-based mRNA enrichment, mRNA fragmentation and complementary DNA (cDNA) synthesis. Sequencing of cDNA libraries was performed in the Illumina NovaSeq platform with 2× 150–base pair (bp) read length. Reads were aligned to the *Mus musculus* GRCm38 reference genome available on ENSEMBL with STAR aligner v.2.5.2b.

#### Bioinformatics data analysis

DESeq2 was used for comparison of gene expression levels between different sample groups. *P* values and log2 fold changes were calculated using a Wald test. Genes with adjusted *P* values (*Padj*) of <0.05 were referred as DEGs. DeconV was used to deconvolute RNA-seq data and infer proportions of DRG subtypes in WT and *Pten^Het^* [39]. DeconV consists of a reference model and a deconvolution model. The reference model learns parameters from single-cell reference, which the deconvolution model uses to infer optimal cell type composition of a bulk sample. We used the adult dataset from Sharma et al. [38] as a single-cell reference and we compiled bulk samples from Accession number GEO: GSE131230, and GSE162263 [64, 65] as the bulk counterpart of the reference sample. DeconV was executed with scripts based on examples provided by the developers. Briefly, we fitted the bulk reference model using the parameters of the single cell reference to obtain a probabilistic model consisting of a discrete distribution (zero inflated negative-binomial) with cell-type specific parameters for single-cell gene counts. Once the reference model was fitted, the deconvolution model was used to translate expression to real bulk gene expression. Based on the combination of parameters from the reference and deconvolution models, we generated a reference file which was then used as a template to infer subtype proportions in our bulk RNA-seq dataset. Default parameters were used for fitting, modeling, and training of the algorithm.

#### Tissue processing and immunolabeling

All immunolabeling was performed on cryosections following standard protocols [54]. Whole mouse embryos, from litters ranging from E9.5 to E14.5, were fixed overnight at 4°C in 4% paraformaldehyde with gentle agitation. For analysis of P21 DRGs, intact spinal columns were first dissected and fixed overnight at 4°C in 4% paraformaldehyde. The next day, spinal columns were washed in PBS for 30 minutes at room temperature, and spinal cords and attached DRGs were dissected from the spinal column for further processing. All tissue was cryoprotected by incubating in 15% sucrose in PBS for 1 hour, followed by 20% sucrose in PBS overnight at 4°C with gentle agitation. Samples were mounted in OCT and rapidly frozen in methylbutane on dry ice. 20μm consecutive sections were cut on a Leica CM3050S cryostat, mounted on slides, and allowed to dry for 2 hours at room temperature.

For immunolabeling, cryosections mounted on slides were washed 3 times for 5 minutes in PBS at room temperature and incubated for 1 hour in blocking solution (5% v/v Normal Donkey Serum, 5% v/v DMSO, 0.25% v/v Triton X-100 in PBS). Slides were then incubated in primary antibodies diluted in blocking solution overnight at 4°C in a humidified staining box. Concentrations for primary antibodies are listed on Table 1.

**Table 1:** Antibodies used.

| Antibody | Species | Supplier | Cat # | RRID | Concentration |
| --- | --- | --- | --- | --- | --- |
| **Primary Antibodies** |  |  |  |  |  |
| Calbindin | Rabbit | Swant | CB 38 | AB_10000340 | 1:1000 |
| Tyrosine<br>Hydroxylase<br>(TH) | Rabbit | Millipore | AB152 | AB_390204 | 1:1000 |
| TrkA | Goat | Fisher (R&D<br>Systems) | AF1056 | AB_2283049 | 1:100 |
| TrkB | Goat | Fisher (R&D<br>Systems) | AF1494 | AB_2155264 | 1:100 |
| TrkC | Goat | Fisher (R&D<br>Systems) | AF1404 | AB_2155412 | 1:100 |
| Casp3 | Rabbit | Cell Signaling | 9661S | AB_2341188 | 1:500 |
| Parvalbumin | Rabbit | Swant | PVG213 | AB_2721207 | 1:1000 |
| Isl1/2 | Mouse<br>Igg2b | DHSB | 39.4D5 | AB_2314683 | 1:250 |
| GFP | Chicken | Abcam | ab13970 | AB_300798 | 1:1000 |
| **Secondary Antibodies** |  |  |  |  |  |
| Anti-Rabbit<br>IgG 488 | Donkey | Thermo<br>Fisher<br>Scientific | A-21206 |  | 1:500 |
| Anti-Rabbit<br>IgG 546 | Donkey | ThermoFisher | A10040 |  | 1:500 |
| Anti-Rabbit<br>IgG 647 | Donkey | ThermoFisher | A-31573 |  | 1:500 |
| Anti-Mouse<br>Igg2b 488 | Goat | ThermoFisher | A-21141 |  | 1:500 |
| Anti-Chicken<br>IgY 488 | Donkey | Jackson<br>Immuno<br>Research | NC0215979 |  | 1:500 |
| Anti-Goat<br>647 | Donkey | ThermoFisher | A-21447 |  | 1:500 |

Slides were then washed three times for 5 minutes in PBS and incubated for 2 hours at room temperature in secondary antibodies diluted at 1:500 in blocking solution. EdU incorporation was detected using Click-iT™ EdU Cell Proliferation Kit for Imaging, Alexa Fluor™ 647 (Thermo Fisher Scientific, Cat#C10340). Slides were incubated in EdU detection solution (100 μM Tris pH7.5, 4 mM CuSO4, 5 μM sulfo-Cy5 azide, 100 mM sodium ascorbate) for 30 min at RT, protected from light. Slides were washed 3 times for 10 minutes in PBS, with 1:5000 Hoechst added to the second wash to label nuclei. Slides were then coverslipped with Fluoromount -G and imaged.

#### Proliferation assays

A single intraperitoneal (IP) injection of EdU in sterile PBS (10µg EdU/g body weight) was given to pregnant dams on one day between E9.5 and E14.5. Two hours after injection, embryos were collected and fixed as described.

#### Cell counts and statistical analysis

To quantify immunoreactive DRG neurons, level matched DRGs from lumbar segments were serially sectioned. For analysis of total number of neurons, sections of entire DRGs were counted. For population proportions, cell proliferation, and cell death assays, 10 consecutive sections of level matched DRGs were counted. Sections were co-immunostained with antibodies against specific cell makers and Isl1/2. The proportions of each population were calculated by dividing the number of marker-positive cells over the number of total neurons in serial sections from level-matched DRGs of control and mutant mice. Values were averaged for each experiment, with a minimum of 3 animals of mixed sex for each genotype at each timepoint and condition across different litters. The average ± SEM is reported in the figures. Statistical analysis for each experiment was performed by Student’s t-test. P-values are reported in figure legends and results section.

---

**Supplemental Figure 1:**
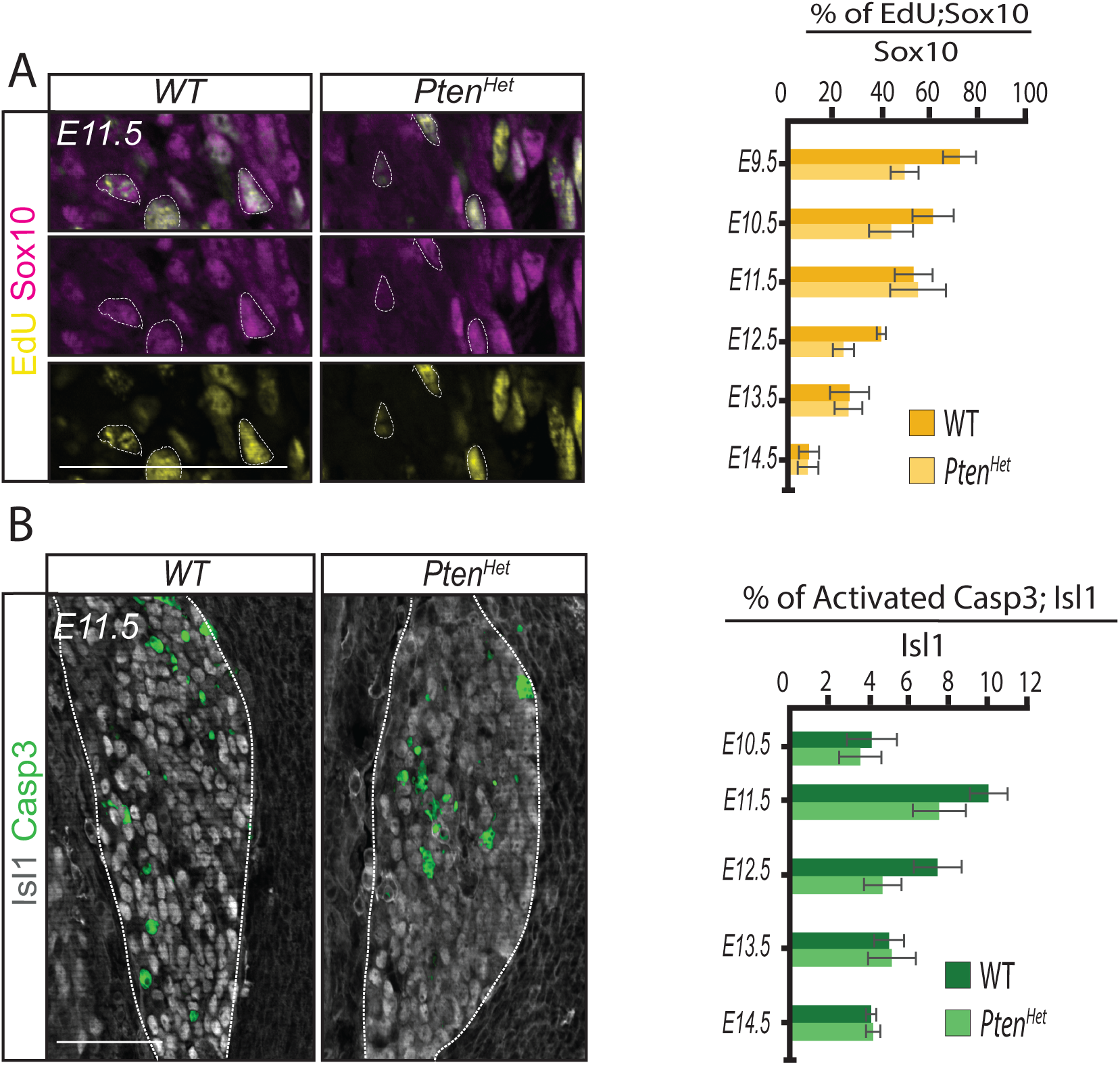
Proliferation and developmental cell death are normal in *Pten^Het^* mutants. **A)** Representative images of E11.5 DRGs from WT and *Pten^Het^* littermates birth-dated with EdU and labeled with Sox10 immunostaining (left panels) (Scale 100 µm). Quantification (right panel) shows percentage of Sox10^+^ progenitors undergoing proliferation at each developmental time point (E9.5-E14.5). At all ages quantified, there is no significant difference in the percentage of EdU^+^;Sox10^+^ precursors in *Pten^Het^* compared to WT controls (n=3, E9.5: p=0.055; E10.5: p=0.11; E11.5: p=0.45; E12.5: p=0.06; E13.5: p=0.484; E14.5: p=0.481, at E14.5; Student’s t-test). **B)** Representative images from E11.5 DRGs of WT and *Pten^Het^* littermates showing activated Caspase 3 (Casp3; green) and Islet 1/2 immunoreactivity (left panels) (Scale 100 µm). Quantification (right panel) shows percentage of Isl1/2^+^ neurons co-labeled with activated Caspase 3. At all ages quantified (E10.5-E14.5), there is no significant difference in the percentage of activated Casp3^+^;Islet1^+^ neurons in *Pten^Het^* DRGs compared to WT controls (n=3, E10.5: p=0.54; E11.5: p=0.16; at E12.5: p=0.069; at E13.5: p=0.91; at E14.5: p=0.84, Student’s t-test).

